# The quiescent X, the replicative Y and the Autosomes

**DOI:** 10.1101/351288

**Authors:** Guillaume Achaz, Serge Gangloff, Benoit Arcangioli

## Abstract

From the analysis of the mutation spectrum in the 2,504 sequenced human genomes from the 1000 genomes project (phase 3), we show that sexual chromosomes (X and Y) exhibit a different proportion of indel mutations than autosomes (A), ranking them X>A>Y. We further show that X chromosomes exhibit a higher ratio of deletion/insertion when compared to autosomes. This simple pattern shows that the recent report that non-dividing quiescent yeast cells accumulate relatively more indels (and particularly deletions) than replicating ones also applies to metazoan cells, including humans. Indeed, the X chromosomes display more indels than the autosomes, having spent more time in quiescent oocytes, whereas the Y chromosomes are solely present in the replicating spermatocytes. From the proportion of indels, we have inferred that *de novo* mutations arising in the maternal lineage are twice more likely to be indels than mutations from the paternal lineage. Our observation, consistent with a recent trio analysis of the spectrum of mutations inherited from the maternal lineage, is likely a major component in our understanding of the origin of anisogamy.

## Introduction

In humans, male and female germ cells are produced before birth and remain quiescent until puberty. At this point, oocytes are released monthly, whereas spermatocytes are continuously dividing to produce spermatozoids (1). Haldane proposed the existence of sex-specific mutation rates, arguing that the frequency of new hemophilic males from non-carrier mothers (hemophilia is X-linked) was very low compared to the expectation of the population frequency, under a mutation-selection equilibrium (2). This deficit of *de novo* mutations in the oocytes was taken as a proof that most mutations occurred in the male lineage. This observation was later connected to the higher evolutionary point mutation rates reported for Y than for X (3–5) and more recently to direct counting of de-novo mutations in humans (6–12), chimpanzees (13) and rodents (14).

However, several lines of evidence both theoretical (15) and from observations on CpG sites (6,11) have suggested that replication-independent mutations accumulate linearly with *absolute* time, regardless of cell division. Because autosome lineages are equally split in both sexes, X chromosome lineages are 2/3 of the time in oocytes and Y lineages are exclusively in spermatozoids, we reasoned that if a quiescence-specific mutational signature exists in humans, it should be revealed by a different mutational spectrum of the X, Y and autosomes.

We have recently shown (16) that the quiescent haploid yeast *Schizosaccharomyces pombe* cells exhibit a distinctive mutational landscape called Chronos: particularly, (i) they accumulate as many indels (insertions or deletions) as SNVs (Single Nucleotide Variants) whereas replicating cells accumulate more SNVs and (ii) they accumulate more deletions than insertions whereas the opposite is observed for dividing cells. The enrichment in indels indicates that DNA lesions also occur during quiescence. Since errors during DNA replication are believed to be responsible for most mutations in many species, we questioned to what extent the replication-independent mutational landscape observed during quiescence in fission yeast also applies to humans.

## Results

To test this idea, we evaluated the distribution of SNVs and indels in human chromosomes. Since indel alleles with fewer than 3 counts are poorly genotyped in autosomes (Supp Fig. 1), we considered the 39.5 million variable sites that host two or more alleles of 3 counts or more each (MAC≥3) in the 2,504 sequenced human genomes from the 1000 genomes project phase 3 (17). For each variable site, we estimated the number of mutations as the number of alleles minus one, leading to a total of 39.8 million mutations for all chromosomes. We also computed the “accessible size” that was sequenced using variants density. To assess the impact of the mutational bias on a longer evolutionary time, we next compared human and chimpanzee genomes, by counting the number of SNVs and indels in a pairwise alignment between both reference genomes. All counts are reported in Supp Table 1.

As population genetics would predict from an effective population size that is 3/4 and 1/4 that of Autosomes for X and Y, respectively, we observed that the density of mutations per base among the variants segregating in humans is ranked Autosomes>X>Y both for indels and SNVs (Fig. 1). Interestingly, at the divergence level while comparing humans and chimpanzees, the degenerating Y chromosome has accumulated more indels and SNVs than any other chromosome (Fig. 1), a pattern that was reported previously (18,19). All differences are highly significant (see Materials and Methods) since counts are typically on the order of millions.

**Figure 1.**
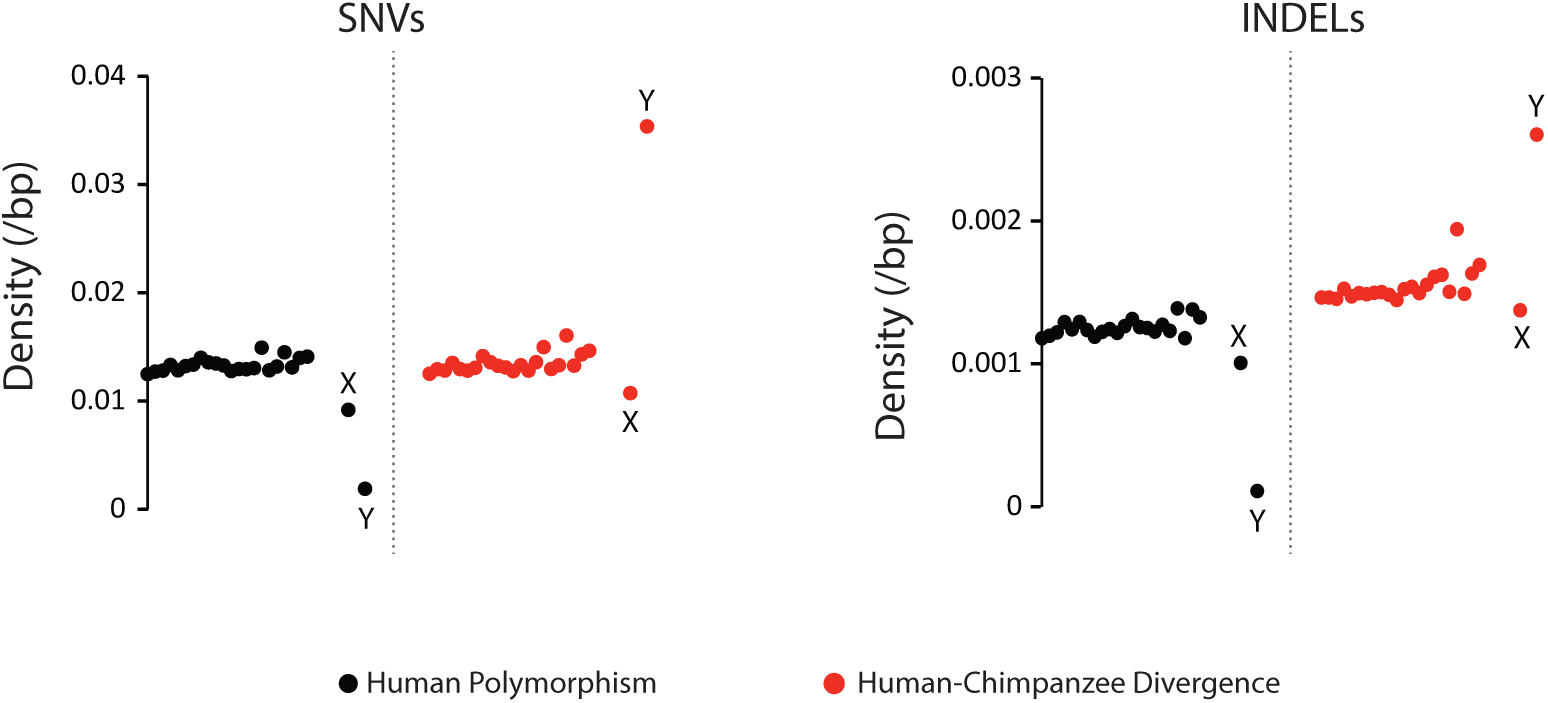
Density of SNVs and indels in X, Y and Autosomes. The density was computed as the number of mutations per bp of the “accessible chromosome” (for segregating polymorphisms) or of the “aligned” chromosome (for human-chimpanzee divergence), which is estimated from the data (see Supplementary Material). Autosomes are ranked by number

We next computed for each chromosome, the fraction of indel mutations among both types of mutations (SNVs and indels). Results show that the fraction of indel mutations is ranked in a strikingly simple pattern: X>Autosomes>Y (0.10>0.09>0.06) (Fig. 2a). Using least squares (see Materials and Methods), we estimated the fraction of indels among *de novo* mutations to be 0.12 in females and 0.06 in males. Finally, we observe that among indels, deletions are even more abundant for the X chromosome (deletion/insertion ratio is 1.9 for X and 1.6 for Autosomes) than for the Autosomes (Fig. 2d). We also noticed a negative correlation between the deletion/insertion ratio and chromosome size (spearman r2=0.58; P=5.8 10-5), but this has not been further investigated yet.

**Figure 2.**
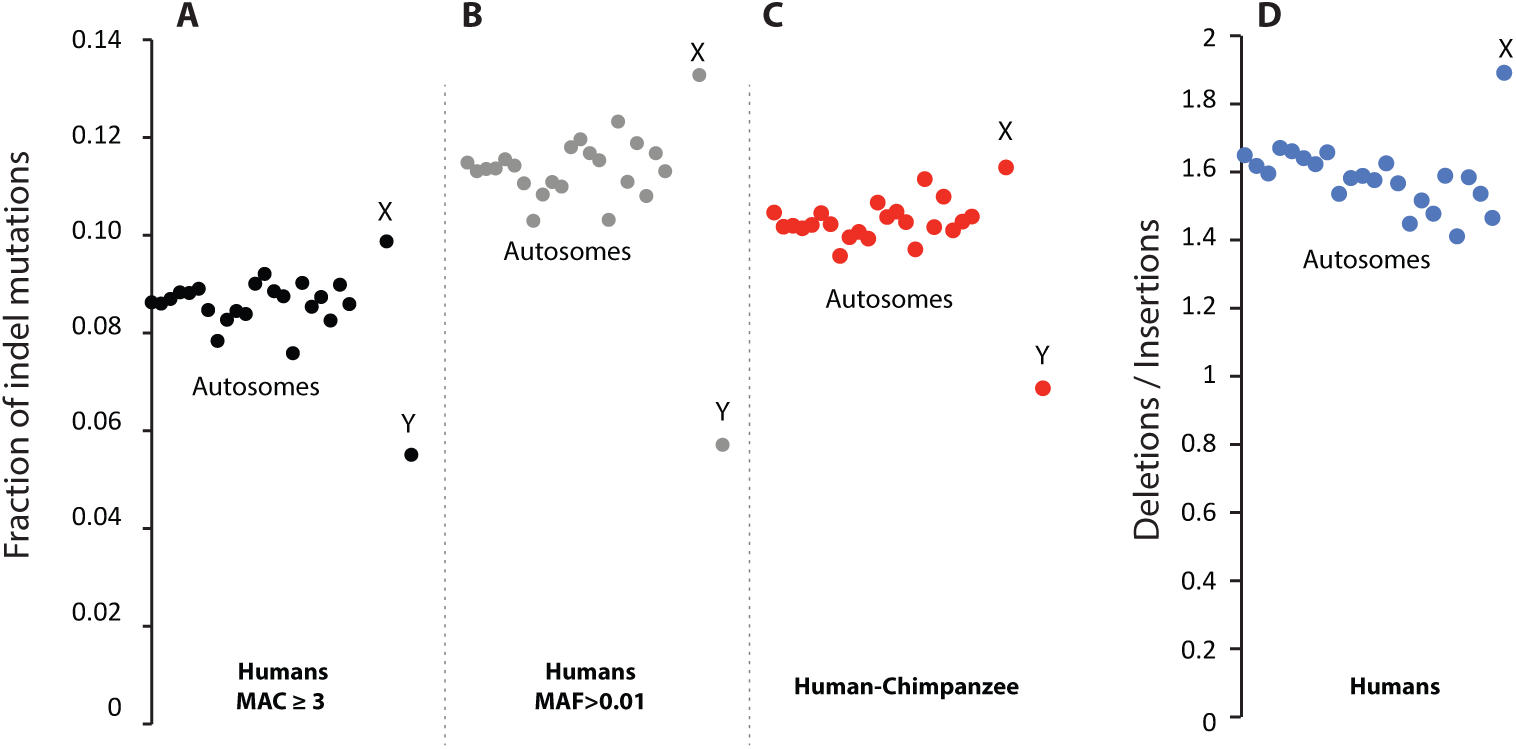
Comparison of Indels among X, Y and Autosomes. **a** We report the fraction of indels segregating in a sample of 2,504 human diploid genomes for the X, the Y and the Autosomes ranked by chromosome number. Only alleles with at least 3 counts (Minor Allele Counts ≥ 3) were considered here because lower frequency indels are poorly genotyped in Autosomes. **b** Variants were further filtered by their frequency (Minor Allele Frequency > 0.01). **c** We provide the same results for the human-chimpanzee comparison. **d** We report the ratio deletion/insertion in all indels that are oriented using the inferred Ancestral Allele. No oriented indel is reported for the Y chromosome or the mitochondria.

At the interspecies level, we observed a very similar pattern (Fig. 2c, 0.11>0.10>0.07). Interestingly, the recent mutations (segregating within humans) contain relatively fewer indels (0.09 on autosomes) than the older ones that have been fixed and detected between human and chimpanzee (0.10 on autosomes). Similarly, the mutations that have reached a 0.01 frequency in the population and are thus more likely to get fixed in humans also exhibit a higher fraction of indels (Fig. 2b). This result supports the view arguing that SNVs are efficiently removed by purifying selection only in the long run (20) likely because they contain many mutations with a small negative fitness impact, the so-called “slightly deleterious” alleles (21–23). Alternatively, one could imagine that indels are not equally efficiently called at all frequencies, thus leading to fewer indels at very low frequencies.

Interestingly, the mitochondrial genome (Supp Table 1) shows a low proportion of indels, reminiscent of the Y chromosome (0.010 of segregating mutations within humans and 0.012 for the human-chimpanzee comparison). Although this low proportion of indels could be due to the high density of functional sequences, it is tempting to postulate that the mitochondria keep dividing and replicating their genome in the quiescent oocytes, therefore masking the imprint of quiescence.

## Discussion

The difference in mutational spectrum between X, Autosomes and Y is unlikely to result from an experimental bias. For instance, sequencing errors or calling bias would likely affect all the “accessible” regions of all chromosomes equally and are very likely absent in mutations detected with a frequency higher than 0.01. Additionally, the fractions of indels for the X chromosome of males or females were analyzed separately and both show a larger value than the autosomes (0.10 for females and 0.11 for males), excluding the possibility that the pattern can be due to an easier X genotyping in males because of haploidy.

Our observations infer that, in humans, there is a relatively higher occurrence of indels in the female gamete lineage. Interestingly, in a 2000 review (24), J. Crow intuitively proposed that female gametes may be responsible for indels and male gametes are responsible for SNVs. Here, we show a striking similarity between yeast, humans and chimpanzee suggesting a conserved trend: quiescence accumulates more indels (deletions). The pattern we report here is in line with recent observations reported for the cause of human mutations. First, the positive correlation between the maternal age and the number of maternally inherited de novo mutations (7) clearly demonstrates that non-replicating mutations accumulate in oocytes. Second, conversion (10) and recombination (9) rates are higher in females than in males; furthermore, the number of recombination events (8) or double-strand-break related mutations (22,23) increases with the mother’s age, demonstrating that DNA breaks occur at a high rate in oocytes and accumulate during quiescence. These breaks are very likely the cause of the non-replicating indels.

Although 80% of the mutations (and more particularly SNVs) originates from the paternal lineage (6), we suspect that the recent progress in indels detection with newer generation technologies will reveal many overlooked indels of maternal origin. It is noteworthy to mention that because quiescence mutations accumulate slowly when compared to replication mutations (16), indels in species with short oocyte quiescence time are expected to be mostly driven by males as it was reported in rodents (14). Ideal test cases could be found in anisogenic species having a long enough development to sexual maturity. Interestingly, comparison of the age of mother and father revealed a complex interplay between the age of the quiescent oocyte and the mutations fixed in the paternal genome (25). In addition, several other factors such as chromatin state (26), transcription levels in testis (27) or even reproductive longevities (28) also alter the maternal and paternal mutational spectrum.

It now remains to be investigated whether the differential contribution of males and females to genome evolution, especially in species with slow development, may have been selected for and whether it relates to the origin of anisogamy.

## Material & Methods

### Computation of the SNVs and indels in the 1,000 human genomes dataset

We retrieved all VCF files from the 1,000 genome phase 3 from ftp.1000genomes.ebi.ac.uk in the /vol1/ftp/release/20130502 directory. Because indels with less than 3 counts were poorly genotyped in autosomes (Supp. Fig. 1A), only alleles with 3 counts or more were considered. A site is assimilated to a SNVs site if all its alleles have length 1. Complex double mutations (mostly in the mitochondrial genome) and Alu insertions were discarded. Others were assimilated to indels. The number of mutations at a given site was computed as the number of alleles minus one.

### Computation of the SNVs and indels in the human-chimpanzee alignment

We retrieved the alignment between the human reference genome (version hg38 that was used in the 1000 genome phase 3) and the reference chimpanzee genome (version panTro4) from http://hgdownload.soe.ucsc.edu/goldenPath/hg38/vsPanTro4/. We considered only the aligned segments where the chromosome number was the same on both species (for chimpanzees, chromosomes annotated 2A and 2B were considered as 2). For the mitochondria genomes, we retrieved the NC_012920.1 (human) and NC_001643.1 (chimpanzee) mitochondrial genomes from NCBI. Both sequences were then globally aligned that resulted in a 16,575 nt alignment. In all alignments, we simply counted the number of point mutations and indels considering all subsequent gap symbols as the same indel event.

### Statistical significance

We tested all the differences using χ^2^ homogeneity tests from counts reported in Supp Table 1 and 2. Autosomes counts were pooled. The differential of indels vs SNVs is highly significant in humans: log10(Pχ^2^) = −1018, df=2, as well as in the human-chimpanzee comparison: log10(Pχ^2^) = −2096; df=2. The difference in deletion vs insertion is also highly significant: log10(Pχ^2^) = - 114, df=1.

### Computation of the chromosome “accessible” sizes

We estimated the genotyped part of each chromosome by summing all regions that include variants that are spaced by less than 1 kb. As reported in the 1,000 genome project (17), this represents on average 90% of the chromosome size, with the exception of the Y where it is 18% of the chromosome. On average, variants are spaced by 35 bp and only ~0.01% are spaced by more than 1 kb. Therefore, counting regions with variants spaced by less than 1 kb is a conservative estimation.

### Least-squares estimations of the sex-specific indel fraction

For Autosomes, X and Y, the observed fraction of indels is given in the vector **I** and the mean time spent in male and female lineages in a matrix **T**:

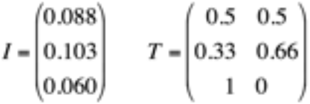

Using standard least squares, we estimated the female and male fraction of indels as (**T T**t)-1**T**t **I**, giving rise to 0.06 for males and 0.12 for females. As a goodness of fit, these values predict that the expected indel fraction in Autosomes, X and Y should be (0.091, 0.101, 0.060). The predicted values are close to those observed in the **I** vector (the difference being (0.003, 0.002, 0.000), see the **I** vector just above).

### Scripts accessibility

All scripts used to parse the data were written in *awk* language and are publicly available at http://doi.org/10.5281/zenodo.2551441.

## Acknowledgments

We would like to thank S. Baulac, F. Débarre, G. Marais, E. Rocha and L. Ségurel for their suggestions on a previous version of the manuscript. We also would like to thank M. Robinson-Rechavi and R. Lanfear who reviewed this manuscript and provided insightful comments. This work was supported by the grants ANR-13-BSV8-0018 and ANR-12-BSV7-0012 Demochips from the Agence Nationale de la Recherche (France) to BA and GA, respectively.

## Conflict of interest disclosure

The authors of this preprint declare that they have no financial conflict of interest with the content of this article. G Achaz is a member of the PCI Evol Biol managing board as well as a PCI Evol Biol recommender

## Supporting Information

**Supplementary Figure 1.**
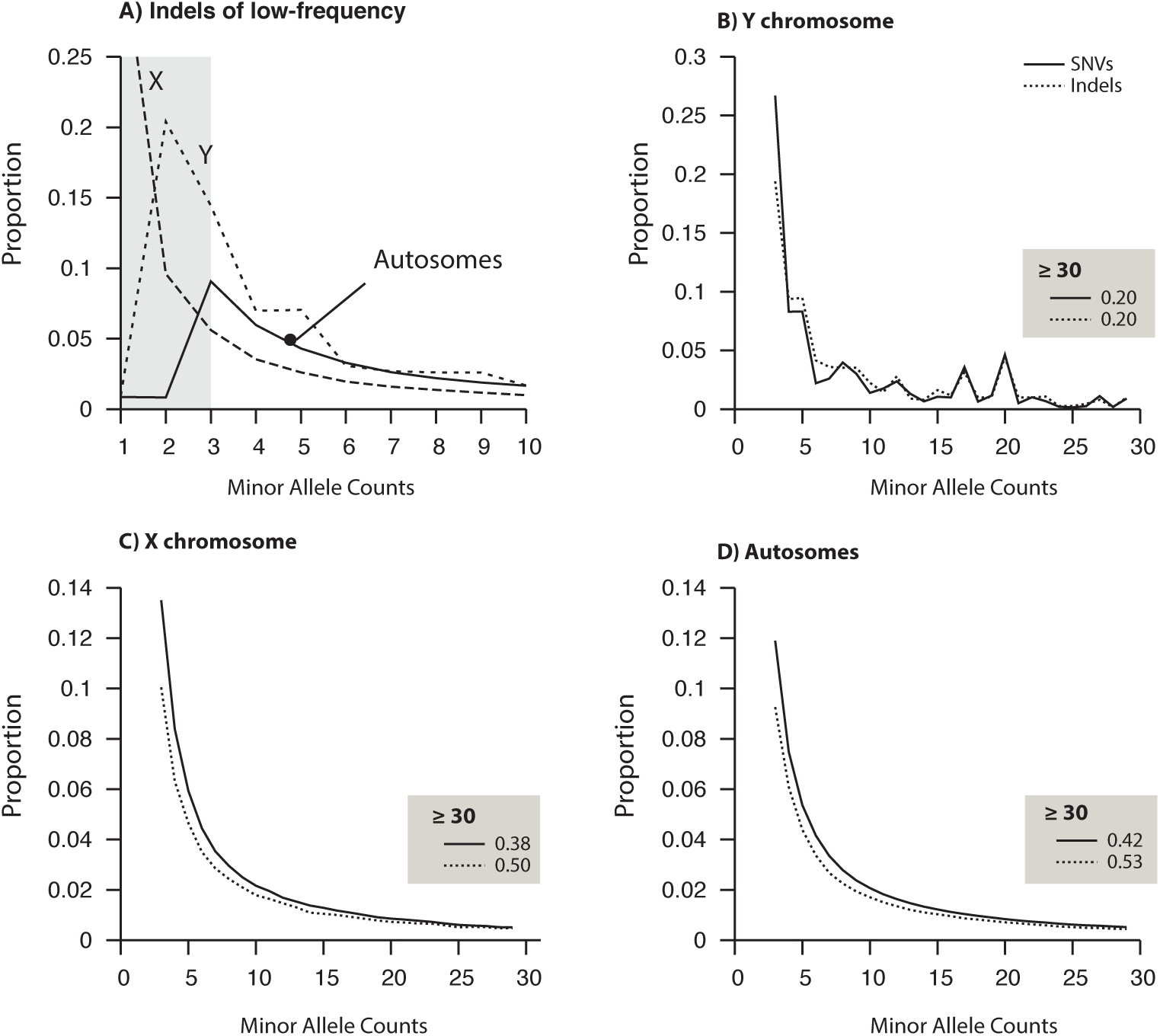
Allele Frequency Spectra (AFS) for SNVs and indels. We report here the distribution of allele frequency (here given by the Minor Allele Count) for bi-allelic variants. A) Normalized AFS for all indels of human chromosomes zoomed on the lowest frequency. B-D) Normalized AFS for SNVs and indels, excluding variants where the minor allele has fewer than 3 counts. Results are given for the Y chromosome (B), the X chromosome (C) and pooled autosomes (D).

**Supplementary Table 1:** total number of SNV and indel mutational events for each human chromosome of the 1,000 human genome project phase 3 (white columns) and in the human-chimpanzee genome alignment (gray columns).

| CHR | Accessible Size | f | #SNV mutations | #indel mutations | Size in the alignments | #SNV mutations | #indel mutations |
| --- | --- | --- | --- | --- | --- | --- | --- |
| 1 | 221,736,733 | 0.89 | 2768960 | 260527 | 214,676,400 | 2,688,535 | 314,370 |
| 2 | 236,052,317 | 0.97 | 3002009 | 281477 | 228,827,157 | 2,959,684 | 335,063 |
| 3 | 194,355,142 | 0.98 | 2489710 | 236234 | 189,206,467 | 2,425,105 | 275,297 |
| 4 | 187,072,586 | 0.98 | 2494816 | 240766 | 180,579,048 | 2,440,890 | 275,447 |
| 5 | 175,575,335 | 0.97 | 2253782 | 217139 | 170,781,708 | 2,211,653 | 251,451 |
| 6 | 166,853,196 | 0.98 | 2207279 | 214962 | 159,651,922 | 2,044,988 | 238,650 |
| 7 | 153,483,682 | 0.10 | 2051182 | 189098 | 148,837,804 | 1,944,712 | 221,314 |
| 8 | 141,740,687 | 0.97 | 1982718 | 168012 | 133,437,499 | 1,887,351 | 199,792 |
| 9 | 113,138,271 | 0.80 | 1535673 | 137968 | 108,925,459 | 1,479,736 | 163,590 |
| 10 | 129,224,414 | 0.95 | 1740547 | 160084 | 122,258,850 | 1,620,335 | 181,258 |
| 11 | 130,592,589 | 0.97 | 1735297 | 158322 | 120,380,559 | 1,577,676 | 174,155 |
| 12 | 130,139,809 | 0.97 | 1661977 | 163920 | 125,126,235 | 1,596,621 | 190,471 |
| 13 | 95,439,646 | 0.83 | 1236299 | 124901 | 85,593,830 | 1,138,577 | 131,714 |
| 14 | 87,931,610 | 0.82 | 1137684 | 110137 | 84,068,261 | 1,077,028 | 125,701 |
| 15 | 79,583,776 | 0.78 | 1038025 | 99186 | 76,611,935 | 1,041,023 | 119,018 |
| 16 | 77,059,256 | 0.85 | 1150388 | 94213 | 73,152,921 | 1,095,981 | 117,689 |
| 17 | 76,927,197 | 0.95 | 987529 | 97678 | 72,619,467 | 941,287 | 117,893 |
| 18 | 74,552,378 | 0.95 | 982968 | 91414 | 72,246,386 | 960,886 | 108,675 |
| 19 | 55,368,119 | 0.94 | 802833 | 76706 | 49,629,257 | 796,857 | 96,378 |
| 20 | 59,328,211 | 0.94 | 777839 | 69758 | 56,442,295 | 749,135 | 84,111 |
| 21 | 34,724,000 | 0.72 | 485412 | 47754 | 33,280,999 | 476,305 | 54,331 |
| 22 | 34,258,169 | 0.67 | 483044 | 45299 | 31,793,980 | 465,627 | 53,803 |
| X | 147,824,448 | 0.95 | 1357783 | 148449 | 125,354,575 | 1,345,122 | 172,364 |
| Y | 10,048,563 | 0.18 | 19022 | 1109 | 16,040,479 | 567,386 | 41,772 |
| MT | 16,545 | 1.00 | 1886 | 20 | 16,575 | 1,451 | 18 |

**Supplementary Table 2:** Annotated Insertions and Deletions for each human chromosome of the 1,000 human genome project phase 3.

| CHR | #insertions | #deletions |
| --- | --- | --- |
| 1 | 30,966 | 50,504 |
| 2 | 34,153 | 54,599 |
| 3 | 29,605 | 46,655 |
| 4 | 29,221 | 48,297 |
| 5 | 26,430 | 43,421 |
| 6 | 26,022 | 42,211 |
| 7 | 22,246 | 35,682 |
| 8 | 20,291 | 33,269 |
| 9 | 16,954 | 25,679 |
| 10 | 19,541 | 30,525 |
| 11 | 19,368 | 30,382 |
| 12 | 19,380 | 30,154 |
| 13 | 15,255 | 24,507 |
| 14 | 13,458 | 20,810 |
| 15 | 12,292 | 17,511 |
| 16 | 10,711 | 16,010 |
| 17 | 11,332 | 16,486 |
| 18 | 11,337 | 17,789 |
| 19 | 7,662 | 10,624 |
| 20 | 8,314 | 13,008 |
| 21 | 6,087 | 9,221 |
| 22 | 5,047 | 7,278 |
| X | 28,109 | 52,822 |

